# The Dynamics of Context-Dependent Space Representations in Auditory Cortex

**DOI:** 10.1101/2023.11.25.568638

**Authors:** Michael H. Myoga, Diana Amaro, Valerija Kello, Matthias Gumbert, Christian Leibold, Michael Pecka, Benedikt Grothe

## Abstract

Auditory space is not mapped onto the receptor surface in the inner ear but results from complex neuronal computations by our brain. However, there is no consensus about the resulting neuronal representation of space in auditory cortex (AC) owing to contradicting reports: While many studies found a tuning preference for sound source positions in the contralateral hemisphere, others have observed an additional population of cells tuned to midline positions or even context-dependent dynamic changes in the tuning of individual cells. A fundamental difference across studies was the animals’ brain state, which might affect AC processing and thus the apparent nature of its spatial code. Yet no study to date investigated spatial tuning of identified AC neurons across brain states. Here, we employed longitudinal two-photon calcium imaging in the AC of mice under distinct states of wakefulness: anesthetized, idle awake, and during involvement in a go/no-go localization task. We find that previously reported differences in coding regimes are directly linked to wakefulness: A strong contralateral tuning bias is present under anesthesia, while pronounced and stable mid-line tuning appeared in awake but idle animals. Intriguingly, in localizing mice, tuning was different again with a large proportion of AC neurons responding to the position of the currently relevant sound-source. These population regimes remained stable across imaging sessions despite the spatial tuning of individual neurons being highly variable. Our findings resolve apparent contradictions in the literature and thus give a dynamical explanation for the multiple space representations in the AC.

## Introduction

The retinotopic organization in primary visual cortex (van Essen, 1985) as well as somatotopy in somatosensory cortex (Penfield and Boldrey, 1937) are organizing principles inherited from the sensory epithelium providing an a priori code of space. In contrast, the auditory system, including auditory cortex (AC), is organized according to tonotopy, i.e., sound frequency. Consequently, the auditory system must compute space based on spectral cues for vertical and binaural differences in level and time of arrival of a sound for the azimuthal sound position. These cues are initially processed at the early synaptic levels of the ascending auditory system (Grothe et al. 2010). It is now generally accepted that – in stark contrast to the analogous neurons of archosaurs (birds and crocodilians) – the azimuthal location in mammals cannot be read out from local, topographically organized labeled line activity of individual neurons (Grothe and Pecka 2014). Rather, binaural responses in the mammalian brainstem and midbrain are largely restricted to the contralateral azimuthal hemifield, independently of whether animals are anesthetized (Yin and Chan, 1990; Grothe et al., 1996; McAlpine et al., 2001, Brand et al., 2002; Pecka et al., 2008) or idle awake (Grothe et al., 1996; Kuwada et al., 2011; Boffi et al,. 2022). Hence, auditory space in mammals is initially encoded via the activity levels of two hemispheric channels (left and right brainstem) by relative (Grothe et al., 2010) and highly dynamic (Boffi et al., 2022) population activity.

How auditory space is represented at the level of the AC, however, remains unclear. Although results from many AC studies are consistent with the two-channel model (Brugge et al., 1994; Stecker et al., 2005; Briley et al., 2013; Yao et al., 2013) recent electrophysiological studies showed additional, in some cases prominent spatial tuning toward the front (Zhou and Wang, 2012). This observation, which has also been corroborated by human psychophysics (Dingle et al., 2010) and EEG studies (Briley et al., 2016), put forth the idea of a third, frontal channel potentially as a starting point for a full representation of all positions at later stages of auditory cortical processing. However, both the mechanistic and anatomical origins of such a third channel, and the conditions under which it is observable is unknown. Moreover, several reports on the dynamics of spatial tuning of AC neurons during behavioral task involvement (Benson et al.,1981; Lee and Middlebrooks, 2011; Salminen et al. 2012; Higgins et al., 2017) contradict rigid channel-based encoding and raise the fundamental question whether, in general, the AC is involved in the maintenance and refinement of sensory space, as found in other sensory systems. Specifically, recent studies indicate the relevance of object-based feature coding in AC, and thus are at odds with traditional ideas of how space information is represented in AC (Wood et al., 2019; Amaro et al., 2021).

A major inconsistency between existing studies is the brain state of the respective subjects during recordings, which ranged from anesthetized, idle awake, to awake & behaving. Since brain state and anesthesia can fundamentally alter processing and coding in sensory cortex (Sabri and Arabzadeh, 2018; Lee et al., 2020), including AC (Ter-Mikaelian et al., 2007), this difference renders it impossible to reconcile the apparently contradictory findings and to identify a coherent scenario of auditory space coding. No study to date investigated spatial tuning of identified AC neurons across brain states.

To overcome this fundamental lack of knowledge and to reveal the context-dependence of spatial coding in AC, we employed longitudinal (several weeks) two-photon calcium imaging of a genetically encoded calcium indicator (GCamp6s) in the AC of CBA/CaJ mice under different conditions and states of wakefulness: anesthetized, awake but idle, and during the performance in a go/no-go localization task. We find that the different coding regimes that had previously been reported are directly linked to differences in behavioral state: A third, mid-line channel only appeared in awake but idle animals while in actively localizing AC, tuning largely centered on the position of the currently relevant sound-source. Therefore, spatial tuning on the level of individual AC neurons is highly variable across recording sessions, while population coding remains stable. Our results thus provide an account for the dynamics of the spatial code in AC, unifying divergent findings in the literature.

## Results

We started by comparing calcium signals in the AC of six mice between anesthetized and awake passively listening states. We imaged in total 1,228 neurons expressing GCamp6s from 12 fields of view (FOVs, 250 × 250 µm) in AC layers II/III over the course of eight sessions per mouse (lasting at least 15 days): four isoflurane-anesthetized sessions were interleaved with four awake (always starting with an anesthetized session). The spatial sensitivity of neurons was probed with amplitude modulated white noise bursts delivered through one of five loudspeakers arranged in a free-field array covering 120 ° of horizontal angular space (30 ° ipsilateral to –90 ° contralateral, Fig. 1a).

**Fig. 1:**
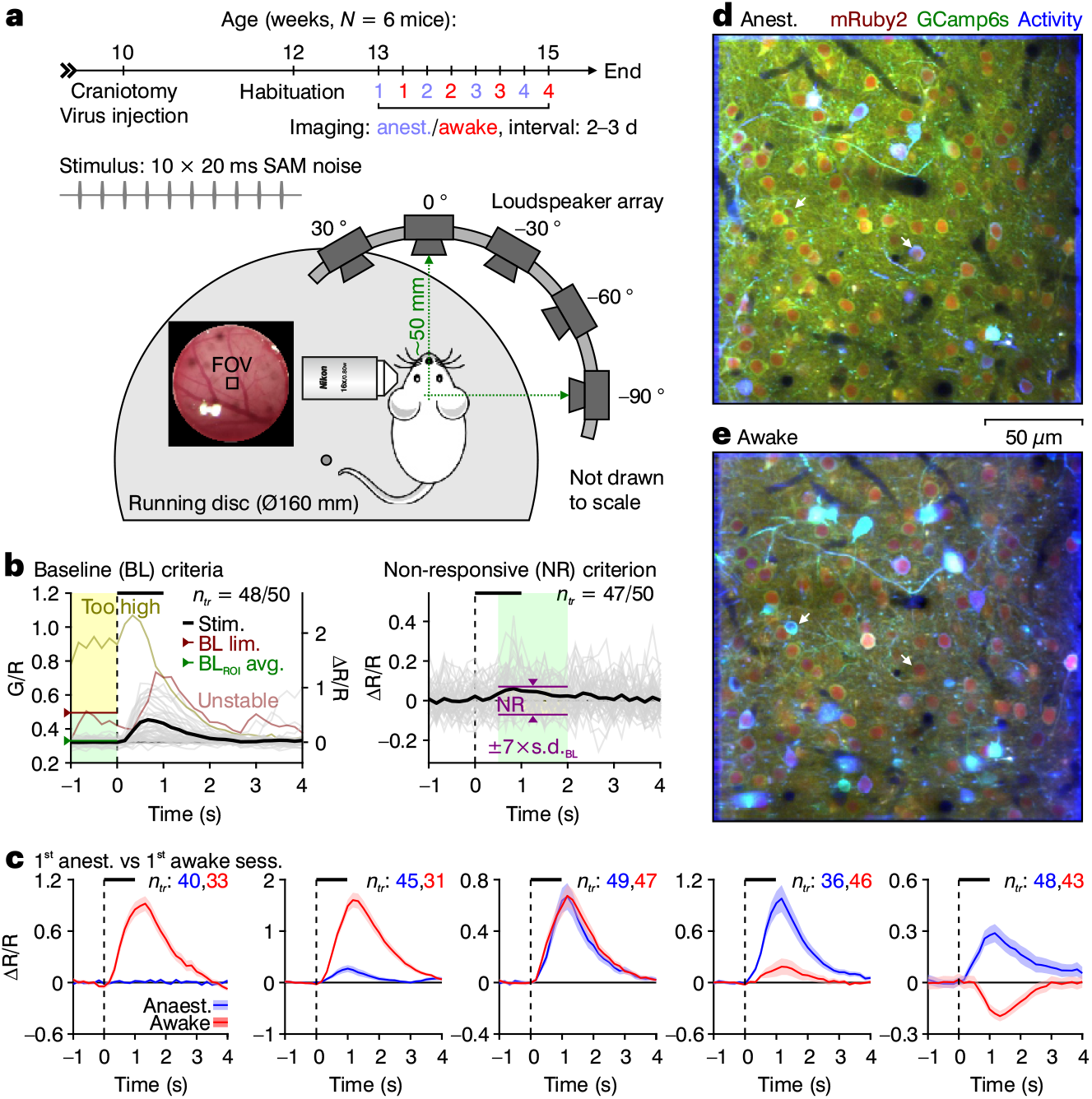
Calcium response amplitude differs between anesthetized and awake states. **a**, Experimental timeline and imaging setup, indicating alternating anesthetized and awake sessions, loudspeaker array and stimulus parameters. **b**, Example fluorescence (G/R, left *y*-axis) traces (gray) from individual regions of interest (ROIs), showing baseline (BL, left) and responsiveness (right) threshold criteria. Individual trials (*n_tr_*: max. 50) were excluded if the baseline was >1.5 times (BL lim., dark red line) the ROI’s session average (BL_ROI_ avg., dark green line) or unstable (exceeded a s.d. threshold). The remaining traces were averaged (black) and baselined to yield ΔR/R values (right *y*-axis), and cells were deemed non-responsive (NR) if the maximum response amplitude was less than seven times the s.d. of the averaged baseline (s.d._BL_, only the 47 accepted trials are plotted). Shaded areas indicate the time window of respective analysis, and green shaded areas represent acceptable values (of baseline G/R or responsiveness). **c**, Averaged accepted calcium traces (± s.e.m., numbers of accepted traces are indicated for each example and state) from example ROIs, showing the diversity of response amplitude differences observed between the first anesthetized and awake sessions. **d**,**e**, Averaged stacks of an example field of view (FOV) in a mouse imaged under anesthesia (**d**) and while awake (**e**). The green and red channels represent averages of the calcium indicator (GCamp6s) and structural marker (mRuby2), respectively, and the blue channel an estimation of neuronal activity based on neighboring pixel correlations over time. White arrows point out example ROIs that were differentially activated between the two wakefulness states.

All quantitative analyses were performed on time-series-extracted regions of interest (ROIs) of visually identified neurons. Because the kinetics of GCamp6s is much slower than the spike rates it indirectly reports, we excluded individual trials without well-defined baseline (see methods for details), as they could contaminate the interpretation of the stimulus-evoked response (Fig. 1b, left). We also determined a responsiveness criterion, where peak calcium responses less than ±7 times the s.d. of the baseline of the averaged accepted trials were deemed non-responsive (NR, Fig. 1b, right).

### Sound-evoked responses are generally higher in the awake AC

We first evaluated the general neuronal responsiveness to sounds between wakefulness states and observed a variety of behaviors. Some neurons were non-responsive in the anesthetized state that became active through the transition (Fig. 1c, left), others showed no distinguishable difference between states (Fig. 1c, center) and others even inverted their response direction (Fig. 1c, right). Despite such diversity, which can also be seen in activity-based pixel maps (Fig. 1d,e, white arrows), sound-evoked responses were on average larger in the awake state (Extended Data Fig. 1), corroborating numerous previous studies (Filipchuk et al 2022; Noda and Takahashi 2015; Gaese and Ostwald, 2001).

### Prominent frontal tuning appears in the awake AC

Relative to the spatial responses of individual neurons, we observed that some neurons exhibited strikingly different spatial tuning between the anesthetized and awake AC, as can be seen in raw calcium traces of an example neuron (Fig. 2a). To quantify the specificity of spatial tuning, we took the peak calcium response to each loudspeaker and generated tuning functions (Supplementary Fig. 1). The best loudspeaker position was determined as the angle of the loudspeaker to which the largest response was elicited. We then performed one-way ANOVAs on each tuning curve, evaluating for statistically significant differences in loudspeaker response amplitude and set specific multiple comparison conditions to categorize best loudspeaker tuning specificity (spec cat, see Methods).

**Fig. 2:**
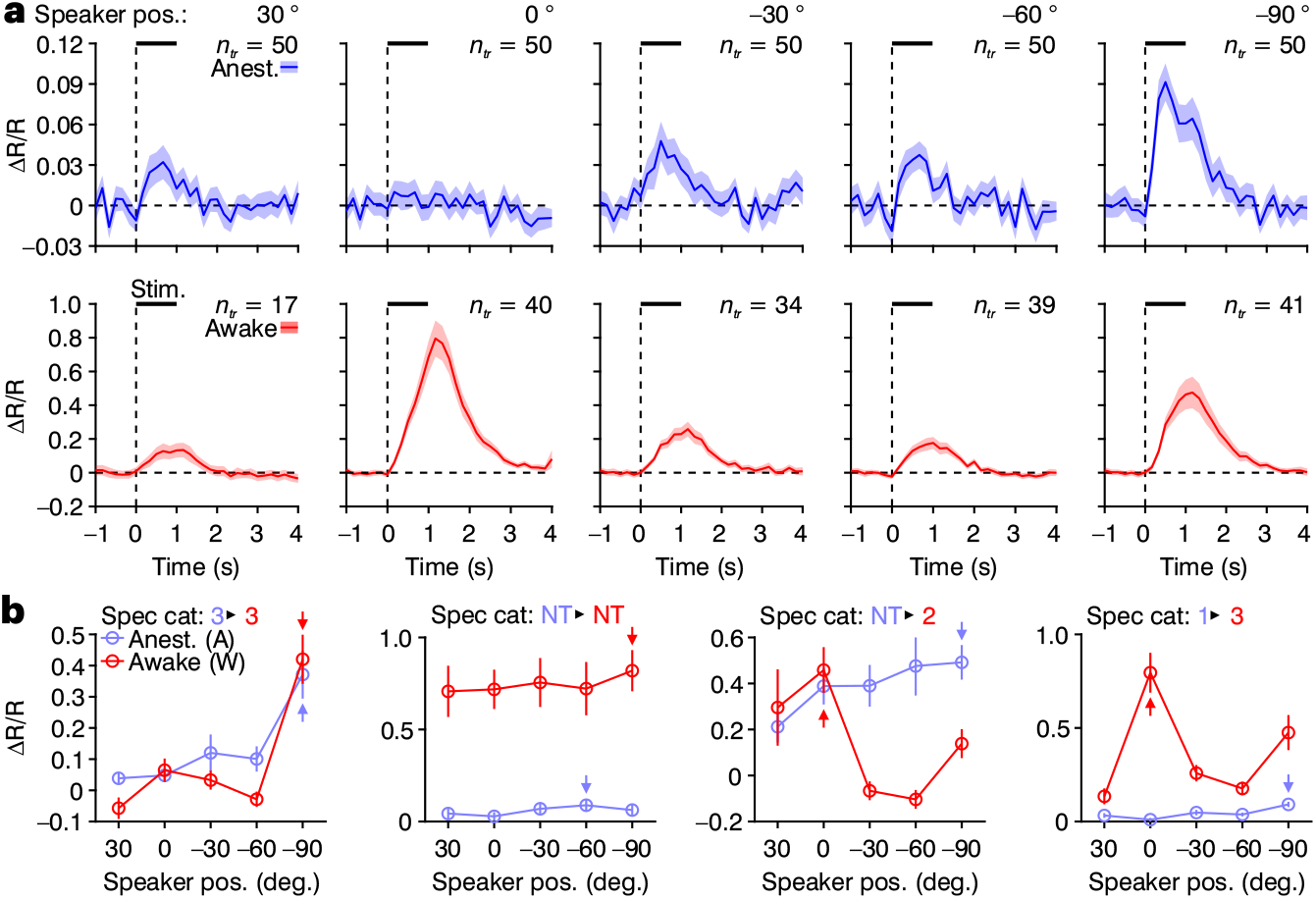
Spatial tuning of individual cells changes between states. **a**, Averaged accepted calcium traces (± s.e.m.) to stimuli from each loudspeaker of an example ROI under anesthesia (top) and while awake (bottom), with the number of accepted trials (*n_tr_*) for each stimulus and state. **b**, Spatial tuning functions of example ROIs in anesthetized (blue) and awake (red) states. Arrows indicate the best loudspeaker and data are mean ± s.e.m..

Evaluating tuning curves of individual cells revealed a diversity of differences between wakefulness states (Fig. 2b). Some cells exhibited no differences in their spatial tuning, either remaining tuned to the same loudspeaker or were non-tuned in both wakefulness states (Fig. 2b, left two panels, respectively). Others became spatially tuned or switched their best loudspeaker (Fig. 2b, right two panels).

To capture the loudspeaker-wise amplitude differences between wakefulness states across the population, we plotted the averaged tuning curves for all tuned cells and the speaker position eliciting the strongest response (best speaker) in both wakefulness states (Fig. 3a). Under anesthesia (blue), spatial tuning of the population was biased to the contralateral side, consistent with numerous previous studies (Stecker and Middlebrooks 2003; Yao et al. 2013; Panniello et al., 2018), and as expected, amplitudes were lower than in awake animals (red). However, the amplitude increases were not equivalent across loudspeakers, with a larger increase at the front of the animal. This difference became more apparent in the ratio of awake to anesthetized amplitudes (Fig. 3b), thus indicating that anesthesia particularly suppresses spatial neuronal responses to the front of the animal.

**Fig. 3:**
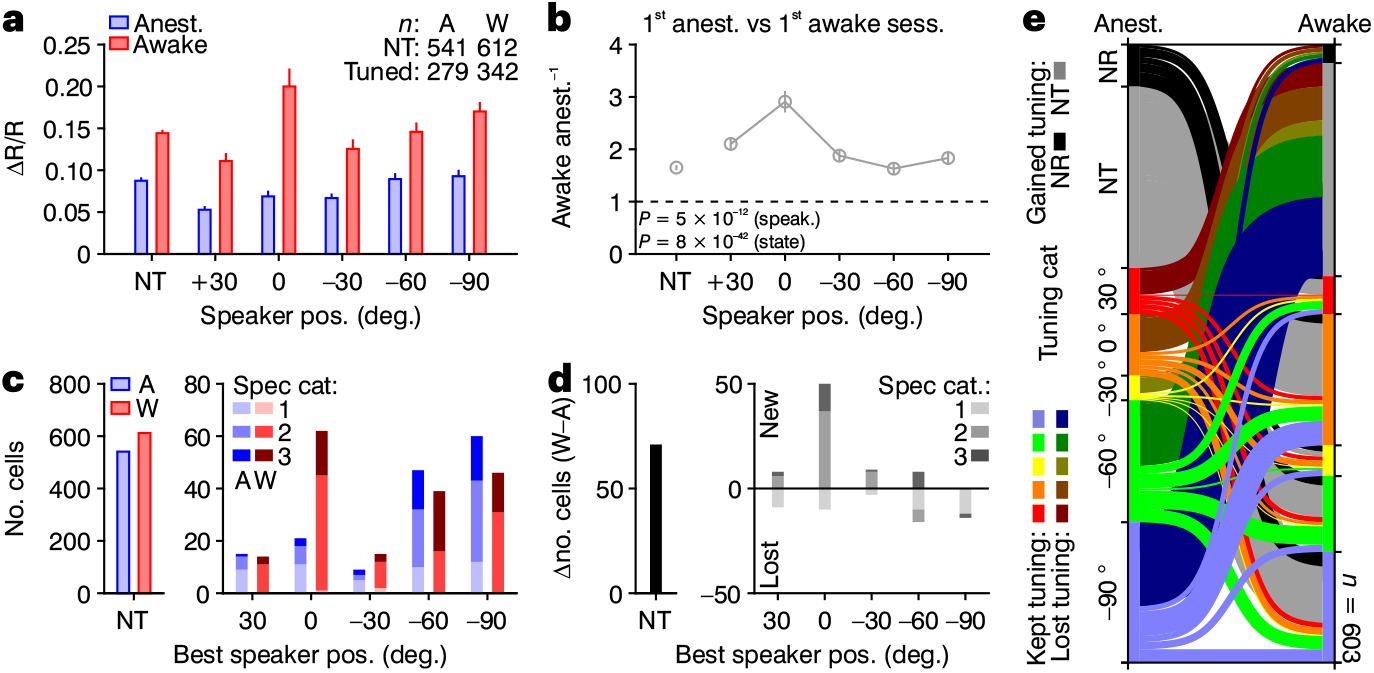
Frontal tuning emerges in the awake AC. **a,** Bar plots of average speaker-wise response amplitudes (± s.e.m.) for tuned cells (lumped spec cats. 1–3 as determined in Supplementary Fig. 1), with maximum amplitudes of non-tuned (NT) cells on the far left. **b**, The ratio of awake to anesthetized response amplitudes of data in **a**. *P* values are the results of a two-way ANOVA test on data shown in **a**, evaluated for loudspeaker and state differences. **c**, Bar plots of the number of cells falling into each spec cat, counted according to best loudspeaker for tuned cells, with non-tuned cells on the left. **d**, Bar plots of the number of new and lost cells of each spec cat in **c** through the first anesthetized to awake transition. **e**, Flow diagrams showing the tuning category switching behavior of cells that were spatially tuned in either state (*n* = 603). The thickness of each line corresponds to the relative number of cells making each transition, and the color code corresponds to tuning categorization in the anesthetized state.

The enhancement of frontal tuning was also reflected in a larger number of cells that became tuned to that loudspeaker position, as can be seen when counting the number of non-tuned cells and all the best loudspeakers of tuned cells in the two wakefulness states (Fig. 3c). In the same animals, substantially more neurons were tuned toward the front in the awake state. Specificity of the tuned neurons was also generally higher in the awake state (note the near absence of spec cat 1 in the awake state). The bias for midline tuned neurons became particularly evident when evaluating the absolute loss and gain of neurons according to best loudspeaker and tuning specificity (Fig. 3d). Thus, a frontal spatial tuning bias appears in the awake state across our data set that is unobservable under anesthesia.

### Frontal tuning in the awake AC arises from diverse tuning preferences under anesthesia

In order to gain insight into the identity of the population of cells that became tuned to the front in the awake state, we generated flow diagrams of all cells that were spatially tuned in at least one wakefulness state to visualize their switching behavior through the first transition, color-coded according to their spatial tuning in the anesthetized state (Fig. 3e, *n* = 603 of 1030 valid ROIs for these sessions). Indeed, some neurons switched their tuning from other loudspeakers to the front, as expected, but the majority of nascent frontally tuned cells were surprisingly non-tuned in the anesthetized state. Interestingly, many other tuned neurons in the anesthetized state also lost their tuning through the transition, which is in accordance with the previous cell counts that revealed slightly more non-tuned cells in the awake condition (Fig. 3c,d, left).

In summary, the gain of frontal tuning in the awake state was reflected in neuronal response amplitudes as well as the number of neurons tuned to the front. The population of cells that gained frontal tuning through the wakefulness state transition was diverse but predominantly non-tuned under anesthesia.

### Frontal tuning in awake animals is prominent and reliable over time in passively listening animals

In order to determine the extent to which frontal tuning in the awake AC in the absence of a behavioral context was stable over time, we next analyzed spatial tuning functions across all sessions (Fig. 4) for valid ROIs over all sessions (Extended Data Fig. 2a,b). Although we observed a general rundown in response amplitudes through the second transition from the anesthetized to the awake state (Fig. 4a and Extended Data Fig. 2c) the general patterns of contralateral bias under anesthesia and frontal bias when awake persisted over the course of the entire experiment (Fig. 4a–d). This can be seen in the response amplitudes (Fig. 4a,b) as well as cell counts (Fig. 4c,d), in accordance with the previous analyses of the first state transition (Fig. 3a-d). Importantly, nearly all imaged FOVs, albeit to different extents, exhibited neurons reliably tuned to the front (Extended Data Fig. 3). Moreover, a generally more specific spatial tuning in the awake AC was reflected in the cumulative observations of tuning specificities over all sessions (Fig. 4e). Thus, the shift towards frontal tuning in the awake AC appears not to be a random drift but rather an engrained feature of the awake but passive AC functional cortical network.

**Fig. 4:**
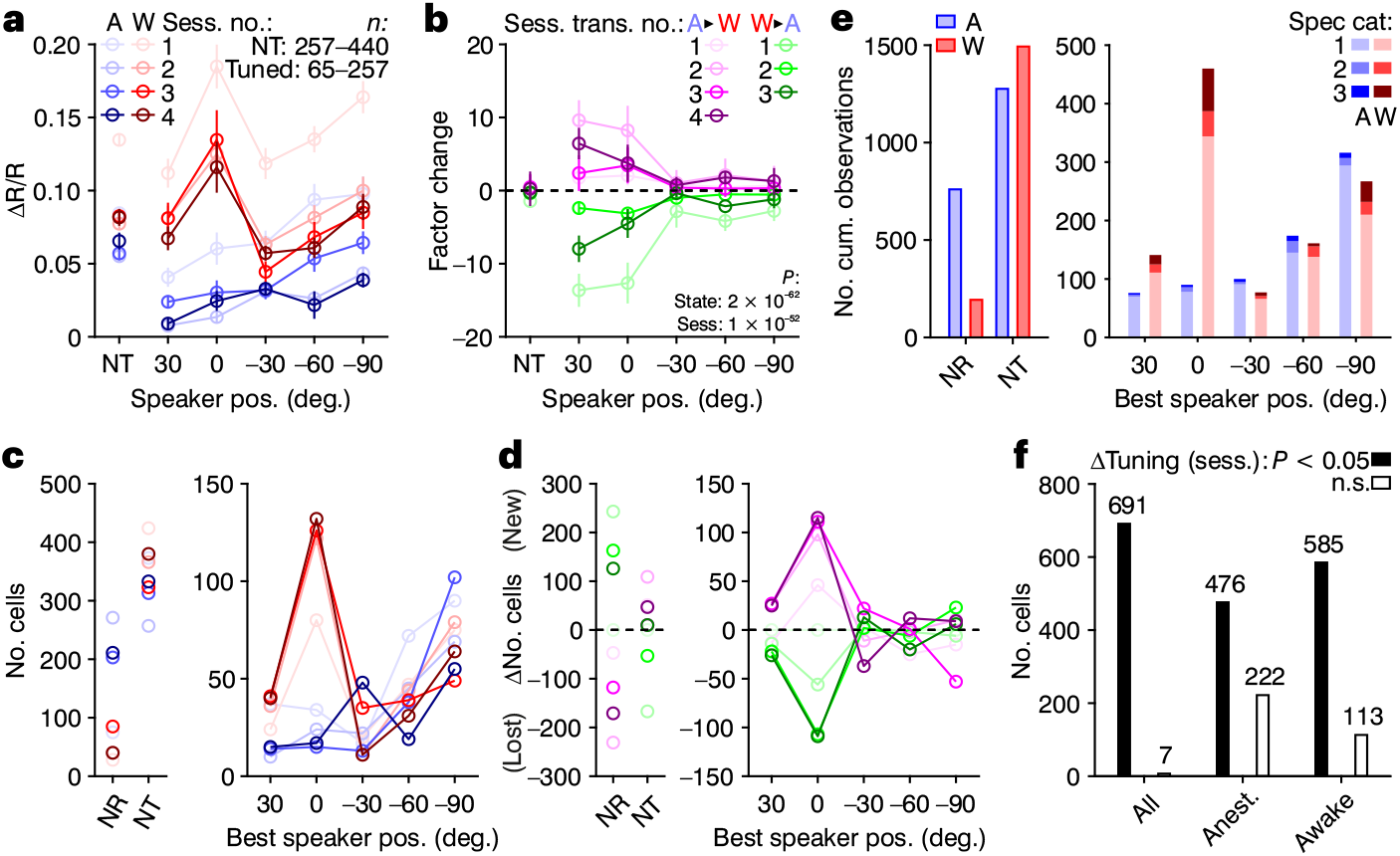
Prominant frontal tuning in awake animals is stable over time. **a**, Average speaker-wise response amplitudes (± s.e.m.) for each anesthetized (blue color scale) and awake (red color scale) session of tuned cells, with maximum amplitudes of non-tuned (NT) cells on the left (as in Fig. 2c). **b**, Factorized change in response amplitude calculated for each state transition, where positive and negative values indicate a relative increase and decrease in amplitude across the transition, respectively. *P* values are the results of three-way ANOVA tests of the data shown in **b**, evaluated for loudspeaker and state differences. **c**, Numbers of tuned cells according to their best loudspeaker for each anesthetized and awake session (as in Fig. 3c, except spec cats 1–3 are lumped together), with those of non-responsive and non-tuned cells on the left. **d**, Change in number of tuned cells according to their best loudspeaker across each state anesthetized-to-awake (magenta color scale) and awake-to-anesthetized (green color scale) transition, with the changes of non-responsive and non-tuned cells on the left (as in Fig. 3d, except spec cats 1–3 are lumped together). **e**, Bar plot of the cumulative number of observations a spec cat was observed, summed over all respective anesthetised (blue) and awake (red) sessions. **f**, Bar plot showing the number of cells that passed (*P* < 0.05, filled bars) or failed (empty bars) a three-way ANOVA test when evaluated for differences in normalized tuning functions across all sessions (left) or two-way ANOVA tests considering only the anesthetized (middle) or awake (right) sessions.

### Frontal tuning in awake animals is maintained largely by the same population of cells

The population of frontally-tuned neurons appearing in the awake AC could consist of different neurons in every session, be always the same neurons, or a combination of the two situations, where some neurons are repeated and others changed between sessions. In the first extreme, mutually exclusive populations of neurons would switch to the front on any given session until the necessary number of frontally tuned neurons is achieved. Looking at the dataset as a whole, one might assume this would be the case since the number of spatially tuned neurons that were common between sessions was low (Extended Data Fig. 2b). Furthermore, only a small fraction of neurons was spatially tuned (to any loudspeaker) on all awake sessions (Extended Data Fig. 2b, right) and therefore the likelihood that those were tuned to the front was low. Moreover, when we performed ANOVAs on normalized tuning curves (from Fig. 4a) and evaluated for statistically significant differences across sessions (Fig. 4f), only seven cells had insignificantly different tuning curves over all sessions (three-way ANOVA). Albeit considerably more across just the anesthetized (222) and awake (113) sessions (two-way ANOVA), these values remain in the minority of the entire population (*n* = 698 valid ROIs). These observations are consistent with the diversity of spatial tuning under anesthesia that comprised the population of frontally tuned neurons in the first awake session (Fig. 3e).

At the other extreme, the same population of neurons would always switch their tuning to the front regardless of what they are tuned to under anesthesia. To address this possibility, we returned to single-ROI analyses. Exploring individual ROIs already indicated some credence to the latter hypothesis, with some units clearly tuned to the front on multiple awake sessions (Fig. 5a, top), while others exhibited rather random switching (Fig. 5a, middle), and others still remained not tuned on all sessions, even though their response amplitude varied from session to session (Fig. 5a, bottom).

**Fig. 5:**
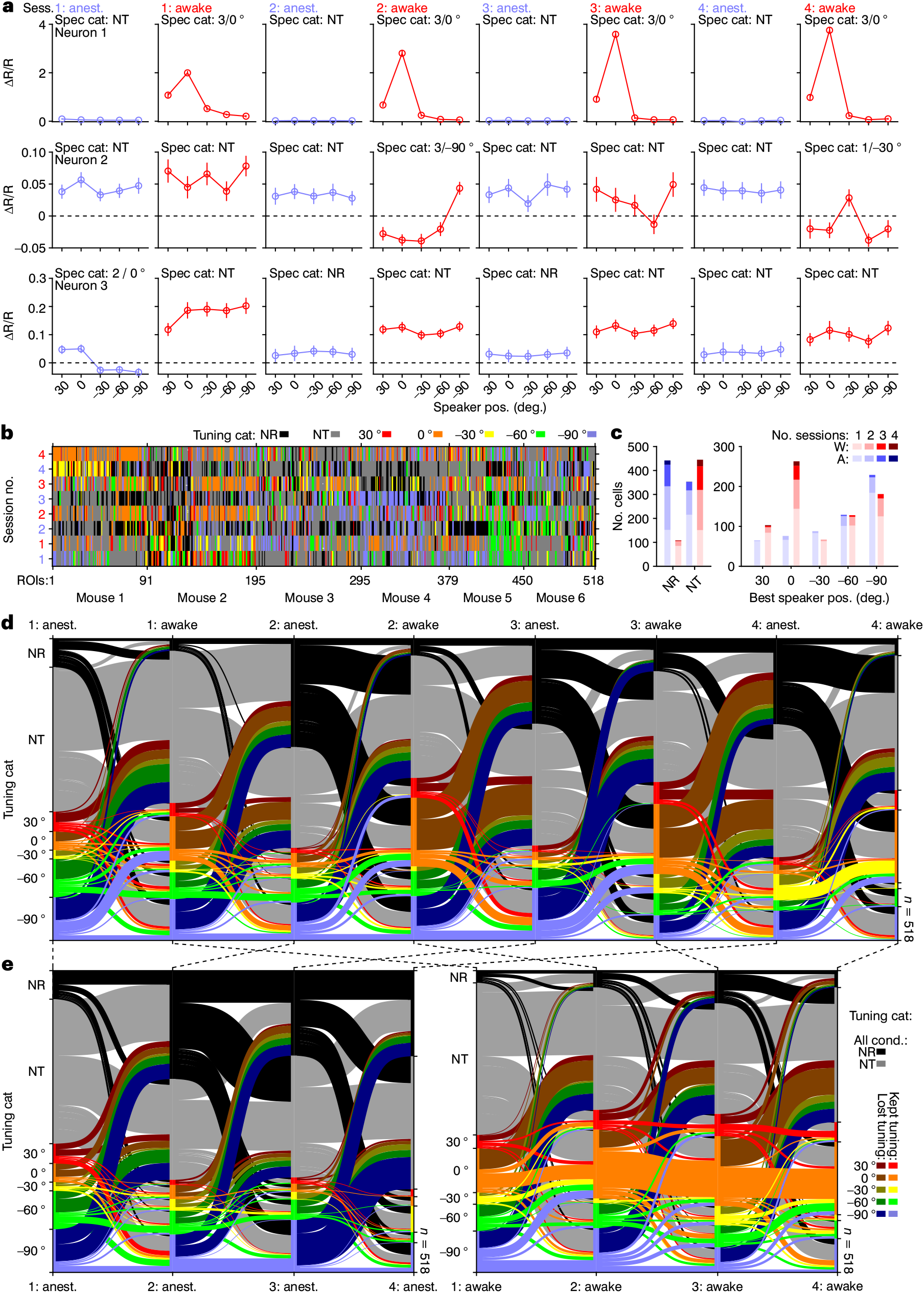
Frontal tuning in awake animals is enforced largely by the same population of cells. **a**, Example tuning functions of 3 cells across all eight sessions (mean ± s.e.m.), with the spec cat (and where applicable, the best loudspeaker) indicated. **b**, Graphical representation of the tuning categorization for all chronically valid ROIs that were tuned on at least one session (*n* = 518 of 698), separated by mouse. Each vertical column represents a single cell’s tuning categorization over all sessions, whereas each row represents the categorization of all cells for any given session or state. **c**, Stacked bar plot showing the number of sessions (awake or anesthetized) a cell fell into a particular tuning category (max. four). **d**, Flow diagrams showing the tuning categorization switching behavior of cells that were spatially tuned on at least one session (*n* = 518) over each state transition in the experiment. The color code represents tuning category in previous (leftward) state. Cells that maintained or lost tuning are in bright and dark colors, respectively, as in Figure 3e. Unlike in Figure 3e, the color codes for non-responsive (black) and non-tuned (gray) cells are the same regardless of whether they maintained a non-tuned status or gained tuning through the transition (each cell in this area would have ventured into the tuned area for at least one session). **e**, Same as in **d**, but flow diagrams show only switching behavior between anesthetized (left) and awake (right) sessions. In all panels, spec. cats. 1–3 are lumped together.

Considering only those neurons that were spatially tuned on at least one session (awake or anesthetized, *n* = 518 of 698 valid ROIs, Extended Data Fig. 2b), we plotted the tuning category of each ROI, separated by mouse, over all eight sessions (Fig. 5b). This inclusive, albeit coarse visualization, gives the impression that at least some neurons were tuned to the front on multiple sessions like the example shown in Figure 5a (top). As a quantification, we next plotted the number of sessions a neuron fell into each tuning category, separated by the anesthetized and awake sessions (Fig. 5c) and found a disproportionate number of neurons tuned to the front in multiple awake sessions. In fact, apart from a few neurons that were never spatially tuned on any awake session (dark red bar segment at NT, Fig. 5c, left) and just two in the 30 ° category, all other neurons falling into the same spatial tuning category on all four awake sessions were tuned to the front (dark red bar segment at 0 °, Fig. 5c, right).

For better visualization, we generated flow diagrams (as in Fig. 3e) across all sessions (Fig. 5d,e). Considering each state transition (Fig. 5d), one can see a random switching behavior, which also seemed largely the case when we evaluated the switching behavior only between anesthetized sessions (Fig. 5e, left). However, considering only the awake-to-awake sessions (Fig. 5e, right) revealed that the majority of neurons maintained frontal tuning across sessions. Thus, prominent and reliable frontal tuning in awake animals is enforced largely by the same population of cells.

### Spatial coding in AC overrepresents behaviorally relevant azimuths

So far, the awake mice were passively listening to the stimuli from the various loudspeakers without any behavioral relevance of speaker location. To determine whether the emergence of a pronounced frontal channel during wakefulness persisted in a sound localization situation, we trained four additional mice in a GO/NOGO paradigm to report sounds played from a particular azimuth while recording neuronal activity of neurons in the AC (Fig. 6a,b). A predetermined rewarded azimuth was kept constant for each mouse, but differed across mice. At the beginning of training, mice generally licked equally often to sounds presented from all azimuths (Fig. 6c). Over time, animals exhibited significantly more licking after a sound was played from the loudspeaker at the rewarded azimuth and abstained from licking to sounds played from other loudspeakers (Fig. 6c). Mistakes were generally made to the neighboring loudspeakers, indicating that the error was most likely reflecting sensory acuity and not difficulties with task design. We then compared neuronal responses to sounds played from different loudspeakers before and after learning (Fig. 6d,e). In all mice, we observed a predominance of responses to the rewarded loudspeaker after learning, which became the largest population of spatially tuned neurons (except for mouse 2, which performed less well and exhibited a tie). As for the comparison between the anesthetized and the passive listening awake condition, this increase was the result of spatial tuning changes of neurons from a variety of neuronal categories as made evident by flow diagrams between sessions before and after learning of the task (Fig. 6e, flow lines are color-coded by the category of the neurons in the session after learning). The population of neurons tuned to the rewarded loudspeaker before learning could have been spatially untuned (Fig. 6d, neuron i), not responsive to sounds (neuron ii), less selective to the rewarded loudspeaker (neuron iii), or it could even have had a non-rewarded loudspeaker as its best loudspeaker, typically the nearest to the rewarded one (neuron iv).

**Fig. 6:**
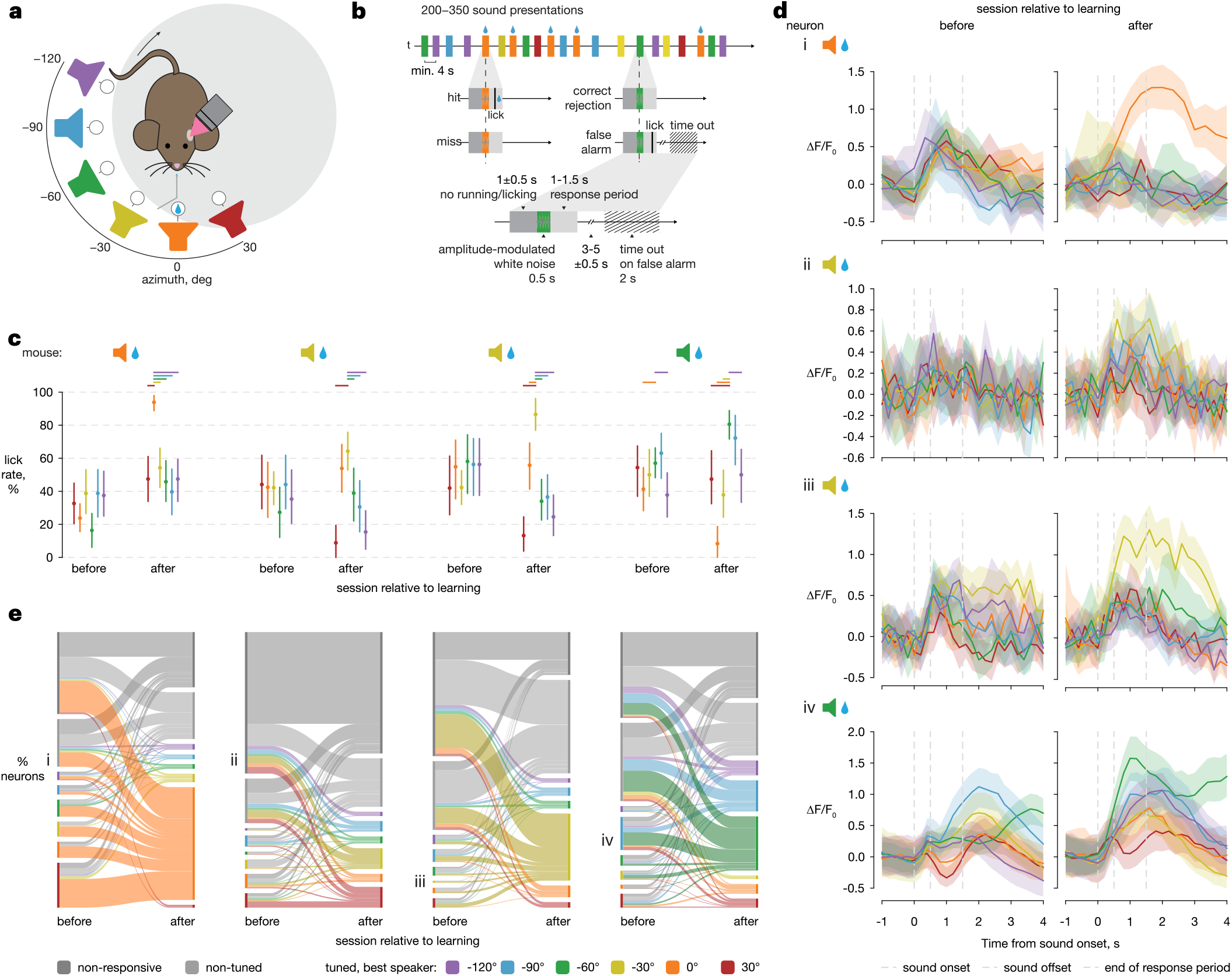
Spatial tuning of single AC neurons reflect spatial behavioral relevance. **a**, Schematic of the behavioral setup depicting a head-fixed mouse running on a disk, a lick spout in front of its mouth and its surrounding array of loudspeakers that span a range of azimuths from 30 ° ipsilateral to 120 ° contralateral in 30 ° steps. For each mouse, the same loudspeaker (in this case the 0° speaker) is always associated with a water reward. **b**, Temporal structure of a behavioral session and possible outcomes for actions following individual sound presentations. The sound presentations from different loudspeakers are represented by colored rectangles following the same color-scheme as in **a**. **c**, Sound localization performance of 4 mice represented by the lick rate in response to the presentation of sounds by each loudspeaker. For each mouse, a session at the beginning of learning and another after task acquisition are depicted. The loudspeaker that was associated with the water reward is shown on top. The error bars correspond to 95% confidence intervals and the horizontal bars indicate to which loudspeakers the animal significantly licked less compared to the rewarded loudspeaker (with Bonferroni correction). **d**, Mean responses of example neurons to each loudspeaker for each mouse in a session before and after learning of the task. The chosen sessions correspond to those in **c**. After learning, all neurons represented were spatially tuned with the rewarded loudspeaker as the best loudspeaker. Before learning, the different neurons had distinct neuronal behaviors: neuron i, non-tuned; neuron ii, non-responsive; neuron iii, best loudspeaker is the rewarded one; neuron iv, best loudspeaker next to the rewarded one. The shaded area corresponds to 95% confidence interval. **e**, Flow diagrams representing the switching behavior of neurons present in both sessions: one before and another after learning (in the same order as in **c** and **d**). Note that color code relates to the “after” learning condition. The colored vertical lines’ lengths are proportional to the percentage of neurons in each session that are in each tuning category - either non-responsive, non-tuned or spatially tuned with a specific best loudspeaker. The lines that connect the before and after sessions represent the percentage of neurons that either change or maintain their state between sessions, color-coded by the state of the neurons in the session after learning. The example neurons depicted in **d** are shown next to the corresponding transition. n_mouse1_ = 253 n_mouse2_ = 384 n_mouse3_ = 432 n_mouse4_ = 450.

The flow diagrams (Fig. 6e) illustrate a significant increase of spatially tuned neurons after learning in all mice (*P* = 0.027, paired *t*-test, *N* = 4 mice, before_m1_ = 46%, after_m1_ = 58%; before_m2_ = 23%, after_m2_ = 30%; before_m3_ = 20%, after_m3_ = 44%; before_m4_ = 32%, after_m4_ = 51%), which, interestingly, was not observable when comparing awake to anesthetized states (Fig. 3c,d). Of the spatially tuned neurons, the percentage having their best loudspeaker as the rewarded one significantly increased (*P* = 0.046, paired *t*-test; before_m1_ = 15%, after_m1_ = 81%; before_m2_ = 17%, after_m2_ = 29%; before_m3_ = 5%, after_m3_ = 64%; before_m4_ = 14%, after_m4_ = 44%). This increase was in part due to the recruitment of new neurons that were non-responsive or non-tuned before learning: these neurons predominantly became spatially tuned to the rewarded loudspeaker. Moreover, of the neurons whose best loudspeaker before learning was the rewarded one and that were still spatially tuned after learning, the majority maintained the same best loudspeaker (100%, 66%, 100% and 50%; total *N*_neurons_ = 12, 3, 3, 8; in the order of the mice). Furthermore, in three out of four mice, neurons that initially were tuned to the loudspeaker next to the rewarded one showed a higher tendency to switch their preference to the rewarded loudspeaker compared to those initially tuned to the further loudspeakers (likelihood of switch to the rewarded loudspeaker from each of the two next loudspeakers or from each of the three further: next_m1_ = 65%, far_m1_ = 46%; next_m2_ = 14%, far_m2_ = 9%; next_m3_ = 27%, far_m3_ = 39%; next_m4_ = 39%, far_m4_ = 11%, *P* = 0.32, paired *t*-test, *N* = 4 mice). In the animal where this trend was absent (m3), the way neurons were recruited for the target location after learning also differed as 50% of those cells were non-responsive before learning (m1 = 28%, m2 = 33%, m4 = 22%). This indicates that various recruitment strategies lead to an increase in the number of neurons responding to the rewarded location.

In conclusion, during a sound localization task, we observed an over-representation of the rewarded azimuth, independently of whether this was in front of the animal or more lateral. The strong response to the frontal speaker seen during the idle awake condition (see above) could not be observed after learning if the frontal position was not the rewarded one. This indicates that the sound localization code in AC is dynamic and dependent on spatial attention and relevance.

## Discussion

Our longitudinal calcium imaging from the AC in mice under three different behavioral states not only questions the traditional ideas of spatial representations in the AC, but also provides a dynamical explanation reconciling existing contradicting reports. Our results indicate a transition of spatial processing from a subcortical encoding strategy of space (hemispheric coding) to a cortical code that appears to be characterized by a third, frontal channel under idle conditions. Yet during active localization behavior, the coding regime is further transformed in context-dependent manner, and we now find the neuronal population to preferentially respond to the spatial position of the currently relevant sound source.

The calcium responses we acquired from AC neurons in anaesthetized mice are consistent with recordings from neurons in the auditory brainstem and the midbrain (inferior colliculus) in anaesthetized (Yin and Chan, 1990; Grothe et al. 1996; McAlpine et al. 2001, Brand et al. 2002; Pecka et al. 2008; Kuwada et al. 2011) as well as awake mammals including monkeys (Grothe et al., 1996; Groh et al. 2003; Werner-Reiss and Groh, 2008; Kuwada et al. 2011; Boffi et al. 2022). The results also agree with those from AC recordings in anaesthetized mammals (Brugge et al., 1994; Stecker et al., 2005; Briley et al., 2013; Yao et al., 2013). Interestingly, we found spatial responses of individual neurons to exhibit large variability across sessions. Despite this large variability at the single-cell level, the responses on the population level were remarkably stable, providing evidence against a labeled line code and in favor of a population code of auditory space in the mammalian auditory system, coherent with recent findings from the auditory midbrain (Boffi et al. 2022, Pecka and Encke 2022).

Our recordings from awake but idle mice show the emergence of numerous responses to the frontal loudspeaker. These frontally-tuned neurons were previously spatially nonspecific or responding to other spatial positions. Interestingly, after acquiring frontal tuning, these neurons exhibited a more stable spatial tuning than neurons tuned to other spatial positions. The finding of frontally tuned neurons in awake mice corresponds well with electrophysiological recordings from awake, also passively listening marmosets (Zhou and Wang, 2012), and EEG recordings from passively listening human subjects (Dingle et al., 2010; Briley et al., 2016). Thus, AC responses to frontal positions are not primate-specific. Noteworthy, recordings from awake marmosets show that cooling of the AC causes, in part, substantial changes in the spatial tuning of a subset of single neurons in the thalamus that project to AC but not in a consistent manner that would constitute a “third channel” (additionally to the two hemispheric channels found in brainstem and midbrain) (Jeschke et al., 2022). This suggests that the existence of frontally tuned neurons in passive hearing mammals is a new feature that arises at the level of the cortex itself. It may result from cortical feedback interactions and relate to the general behavioral importance of objects or events in front of an animal in absence of any currently relevant sound source (position).

In line with this hypothesis, we observed that the prominent response to the frontal speaker could not be observed when mice participated in the GO/NO-GO task, even at the beginning of training. Rather, it appears that, during active listening, when the context of spatial information has fundamentally changed by training the animals to attend to a specific speaker position (in contrast to all other positions that the animal should avoid), neurons tuned towards the presently relevant positions. As a result, we found a clear over-representation of the rewarded azimuthal position, independently of whether this was in front of the animal or more lateral.

The coding of auditory space we find in our experiments follows similar principles as the coding of other stimulus parameters in the AC. Previous studies demonstrated that there is a contextual modulation of neuronal activity in the primary sensory areas (Fritz et al., 2005; Khan et al. 2018; Lee and Middlebrooks, 2011; Corbo et al., 2022; Poort et al. 2022). Compared to passive conditions, when the ferrets perform a tone detection task, neurons of the A1 adapt their spectrotemporal receptive fields by increasing their response to the target frequency (Fritz et al., 2003). Moreover, the same tone can elicit differential neuronal responses depending on the required behavior in the auditory task (Fritz et al., 2005). In the AC of mice, neuronal responses related to behavioral choice in a pure-tone discrimination task have been described (Francis et al. 2022). Recent experiments in the primary visual cortex also revealed a context-depended encoding of drifting gratings when comparing trained behaving mice with trained disengaged or naïve animals (Corbo et al., 2022), and the population encoding of the task context is independent of the representation of the visual stimulus (Hajnal et al., 2023).

In conclusion, our longitudinal recordings demonstrate that the dynamics of spatial tuning follow a specific recruitment logic. Rather than representing space per se, our findings indicate that neuronal activity in AC is flexibly allocated to behaviorally relevant objects in space. Such a re-interpretation of auditory spatial processing at higher levels of the auditory system (AC and downstream) would provide a unifying principle for object identification (Wood et al. 2019, Amaro et al. 2022) or object segregation (Lingner et al. 2018) based on spatial information.

## Methods

### Animals

All animal procedures were approved by the regional animal welfare authority (Regierung von Oberbayern, license no. ROB-55.22532.Vet_02-17-221). In total, six male CBA/CaJ mice (∼10 weeks of age) were used for passive listening experiments and an additional three male and one female (∼12 weeks of age) for the behavioral experiments. Mice were bred and housed in the animal facility of the Max Planck Institute of Biochemistry under a normal 12/12 h light cycle with *ad libitum* access to food and water. For the behavioral experiments, in the training period, the access to water was restricted to the water received while performing the behavioural task and, if necessary, supplemental water was given to ensure that the animals’ weight was kept above 85% of their initial weight.

### Surgical procedures and virus injection

Prior to surgery, animals were anesthetized with an intraperitoneal (IP) injection of a mixture of fentanyl (0.05 mg kg^−1^, Hexal), midazolam (5 mg kg^−1^, Ratiopharm,) and medetomedine hydrochloride (0.5 mg kg^−1^, Orion Pharma). A subcutaneous (SC) injection of carprofen (5 mg kg^−1^, Zoetis) was additionally administered as an analgesic. Depth of anesthesia was determined by the toe-pinch reflex regularly throughout the surgery, and up to an additional 20% of the original anesthesia mixture was injected SC if an animal exhibited the reflex at any time during the operation.

Once fully anesthetized, Isopto-Max eye cream (Alcon Pharma) was applied to prevent corneal drying during the surgery, and animals were placed on a custom-made, heated stereotaxic frame for head bar mounting. The skin of the skull was locally anesthetized with topical application of 10% lidocaine (Xylocaine, Astra Zeneca), and a circular patch of skin (∼1 cm^2^) was removed such that lambda, bregma and bone sutures were exposed. The periosteum was then removed using small dental brushes (Adjustable Precision Applicator Brushes, Parkell), and the skull was further roughened by application of dentin activator (Universal Dentin Activator Gel, Parkell) and carefully etched with a scalpel in a grid-like fashion. The skull surface was then cleaned with sterile cortex buffer (125 mM NaCl, 5 mM KCl, 10 mM glucose, 10 mM HEPES, 2 mM CaCl_2_×2H_2_O, and 2 mM MgSO_4_×7H_2_O) and allowed to dry. Next, a custom-made aluminum head bar (15×3.5×3 mm) was placed on the top of the scull and secured with C&B Metabond dental cement (Parkell). After ∼10 min. of curing time, animals were relocated to a second custom-made, heated stereotaxic frame where the animal could be rigidly fixated via its head bar.

To expose the temporal side of the skull under which the AC is located, the skin around the area was removed. Lidocaine was applied topically to the left temporal muscle, after which the muscle was cut away with scissors. A small craniotomy was then made using a 3 mm disposable biopsy punch (Kai Medical) to expose the target brain area with the dura left intact. The location of the AC was estimated using vascular landmarks according to Joachimsthaler et al. (2014), which determined the location of the injection sites. From this point onward (until resealing the craniotomy), the brain was regularly irrigated with cortex buffer to prevent drying.

For virus injections, borosilicate glass capillaries (cat. no. 1408472, Hilgenberg) were pulled on a P-97 micropipette puller (Sutter Instruments), broken to a tip diameter of Ø40–50 µm, and beveled to an attack angle of ∼35 ° on a custom-made beveller. Finished micropipettes were then front-loaded with a 1:1 dilution (in cortex buffer) of adeno-associated viruses encoding the genetically-encoded calcium indicator GCaMP6s and mRuby2 as a structural marker under the control of the synapsin 1 promoter (pAAV-hSyn1-mRuby2-GSG-P2A-GCaMP6s-WPRE-pA, cat. no. 50942-AAV1, Addgene) and slowly lowered to ∼350 µm below the surface of the exposed brain with an HO-10 hydraulic micromanipulator (Narishige). After 60 s settling time, virus was injected with a Toohey Spritzer Pressure System 2e (Toohey Corporation) for 20–40 ms at 20–40 PSI (to achieve ∼10 nl per injection) every 5 s until a total of ∼250 nl was injected. After one additional minute of recovery time, the micropipette was lifted out of the brain in steps of 100 µm every 60 s. This procedure was repeated to achieve a total of four injection sites (spaced ∼500 µm apart) per animal.

Following the injections, a Ø3 mm circular glass coverslip (no. 1, cat. no. 41001103, Glasswarenfabrik Karl Hecht) was placed over the craniotomy and sealed with Ultra Gel cyanoacrylate adhesive (Pattex). Both the head bar and cover glass were then further stabilized using Paladur dental cement (Heraeus Kulzer), and any remaining loose skin was fixated with Histoacryl tissue adhesive (B. Braun).

At the end of the surgical procedure, the general anesthesia was reversed with an IP injection of Naloxon (1.2 mg kg^−1^, Ratiopharm), Flumazenil (0.5 mg kg^−1^, Hexal), and Atipamezole (2.5 mg kg^−1^, Orion Pharma). Animals were kept under a heat lamp until they exhibited full recovery from anesthesia before being transferred back to the animal facility. The recovery of the animal was monitored for three consecutive days following the surgery, on the sixth post-surgery day, and weekly thereafter. Carpofen (5 mg kg^−1^) was given SC on the two days following the surgery.

### Imaging setup

The imaging setup consisted of a rotatable Bergamo 2 Series benchtop two-photon microscope (Thorlabs), equipped with an 8 kHz resonant scanner, powered by a MaiTai eHP DS tunable Ti:Sapphire laser (Spectra-Physics), and modulated by a Model 350-80 Pockels Cell (Conoptics). The entire microscope chassis was housed in a custom-built, sound-attenuating chamber made out of an aluminum rail-based frame, 5 mm thick ABS panels and foam. In order to image the left AC, the microscope was rotated to 55–65 °, and a 16× 0.8 NA objective (Nikon) was used for all imaging experiments.

The head fixation setup consisted of a custom-made adjustable head bar holder attached to a Ø1.5” Thorlabs stainless steel post system and a Ø160 mm running disc positioned below the head bar holder. Running activity was measured by an MA3 miniature absolute magnetic shaft encoder (Pewatron).

### Stimuli and sound presentation

Amplitude modulated white noise bursts were generated using custom-written functions in MATLAB (version R2018a, Mathworks) at a sampling rate of 192 kHz.

For passive listening experiments, one stimulus consisted of ten 20 ms white noise bursts with 5 ms cosine ramps, played at 10 Hz. Stimuli were analogized on a FireFace UFX II DAC (RME), amplified on an AVR 347 multichannel amplifier (Harman/Kardon) and delivered through one of five CDMG15008-03A dynamic loudspeakers (CUI Devices).

For passive listening experiments, the five loudspeakers were mounted in a custom-made semi-circular aluminum hoop, spaced ∼5 cm away from the center of the animal’s head and positioned 30° apart to achieve an angular spatial extent of 30 ° (ipsilateral) to –90 ° (contralateral). Due to the necessary positioning of the microscope objective to the left of the mouse, further ipsilateral loudspeaker positions were not possible. During experiments, stimuli were presented every four seconds to one loudspeaker in a pseudorandomized fashion, and a total of 50 trials at 4 s inter-trial-interval (ITI) were performed per loudspeaker and stimulus (white or band-passed noise).

For the sound localization task, an additional loudspeaker was added at the azimuth of –120°, and stimuli were presented from the six loudspeakers at 60 dB SPL. The amplitude of the sound was roved ± 5 dB in the training sessions to prevent that the animal could potentially use this cue to solve the task. During imaging sessions, the amplitude was not roved and the noise was frozen.

Each stimulus used during the training phase where the animals learned to associate reward and acoustic stimuli lasted 460 ms and consisted of five 60 ms pure tones of either 3 kHz or 12 kHz with a 10 ms on- and offset cosine ramp that were repeated at 10 Hz. Each stimulus used during the sound localization task lasted 460 ms and consisted of five 60 ms white noise bursts with a 10 ms on- and offset cosine ramp repeated at 10 Hz.

Loudspeakers were calibrated via impulse response to a flat frequency response between 3–70 kHz and a sound pressure level of 75 dB SPL (RMS). Here, a 10 s noise was played through each speaker and measured with a Type 4393 ¼” free-field transducer connected to a Type 2670 preamplifier (Brüel & Kjaer), placed 5 cm away from the loudspeaker, and the resulting transfer function was corrected for phase and amplitude in the Fourier domain using custom-written MATLAB scripts. To calibrate SPL, the recorded signal from a 94 dB SPL 1 kHz pure tone of an SLC 1356 Sound Level Meter Calibrator (RS Components) was used as a reference amplitude. Sounds were digitally attenuated before being sent to the power amplifier.

### Animal handling and anesthesia during imaging

Two weeks after the surgery, mice were handled daily for up to one week to habituate them to head-fixation and the experimental setup (Fig. 1a). First, animals were exposed to the experimenter’s hand. Once an animal crawled on the experimenter’s hand willfully, it was taken to the setup and given several minutes to explore. Animals were then habituated to head fixation for ten minutes while sounds were presented occasionally to accustom them to the stimulus presentation. For awake experiments, mice were allowed to run freely at all times while head-fixed.

For anesthetized experiments, isoflurane was administered in oxygen with a Vapor 19.3 vaporizer (Dräger) and delivered to the animal with an OC-SFM-KIT rodent facemask (WPI). After head and mask fixation, an initial dose of 5% was administered at a flow rate of 2 l min.^−1^ for two minutes. Thereafter, the concentration was reduced to 1% and the flow rate to 0.75 l min.^−1^ for the remainder of the experiment. Isoptomax eye cream was applied directly after the induction of anesthesia, and physiological body temperature was maintained with a heat lamp placed nearby the animal’s body.

Beginning with the start of habituation to the experimental setup, animal wellbeing was again monitored daily until the end of the experiment, when animals were killed by cervical dislocation.

### Longitudinal two-photon calcium imaging

The two-photon microscope was controlled by Scanimage software (Version 4.3, Vidrio Technologies) running on MATLAB (version R2013b). Excitation wavelength was set to 940 nm to excite both GCamp6s and mRuby2. Images were acquired at a depth of ∼200 µm below the pial surface, which required an average laser power of 25–40 mW. Once an auditory-responsive region was found, the scan zoom factor was set to achieve a field of view (FOV) of 250×250 µm. Image dimensions were set at 512×512 pixels, achieving a raw frame rate of ∼30 Hz. Two such FOVs per mouse were imaged over the course of the experiment.

The experimental timeline varied slightly from mouse to mouse, but adhered to the following schedule (Fig. 1a). The first imaging session (3 weeks after virus injection) was always anesthetized. Then, experiments were alternated between awake and anesthetized conditions for a total of eight sessions, with an interval of 2–3 days. At each session, extreme care was made to ensure that the same plane was re-imaged, which required iterative adjustment of the microscope and mouse.

During the sound localization task, images were acquired at 15Hz at a depth of ∼150 µm below the pial surface (superficial layers II/III) corresponding to a field of view of 500 x 500 µm with a resolution of 1024×1024 pixels. The same FOV was recorded before and after the animal learned the task (the time interval between the two varied between 9 and 31 days).

### Behavioral training

The training of mice started with the restriction of their water consumption at least two weeks after surgery, decreasing their weight to a minimum of 85% of their initial weight. At first, the animals were habituated to receive their daily water from a 16 Gauge, 3 mm tip-diameter reusable feeding needle (Fine Science Tools), operated using a normally closed pinch valve (NResearch, Inc.), when they were on top of the hand of the experimenter. Once this was consistently achieved, the animals were accustomed to head-fixation in the setup and trained to lick the lick spout located in front of their mouth to activate the automatic delivery of a waterdrop (5-10 µl). As soon as the animals drank consistently more than 40 drops in one session (lasting maximum 15 minutes), the next training phase was introduced. Up to this phase (one to two weeks after beginning of water restriction) there were no sound presentations and the animals were free to run on the running disk. Afterwards, the animals learned to stop running and licking for a time period of 1 ± 0.5 s to hear an amplitude modulated pure tone, after which they could lick the lick spout to activate a waterdrop delivery. This response period was initially long (3s) and was successively shortened to finally last 1-1.5 s. The pure tone was played from one of the six loudspeakers positioned around the animal, whose location was unimportant for the task. This phase allowed the animals to associate water reward with acoustic stimuli, which is important to the final phase (Fig. 6b), when the animals learned to distinguish and report different sound directions. Again here, the animals had to stop running on the disk and licking the lick spout for 1 ± 0.5 s to trigger the presentation of a stimulus: amplitude-modulated white noise bursts from one of the six possible loudspeakers. During the subsequent response period, if the animals licked when the sound was presented from the target loudspeaker, they were rewarded with a waterdrop (hit). However, when they incorrectly licked to a sound played from another loudspeaker (false alarm), a time out was introduced by adding 2 s to the minimum inter-trial interval of 3-5s. If the animal failed to report a sound presentation from the target loudspeaker (miss) or abstained from licking after a sound presentation from the non-target loudspeakers (correct rejection), it received no water reward.

If an equal number of sounds had been played from each loudspeaker, the likelihood of a sound being played from the target loudspeaker would have been very small (17%), which could have resulted in a lack of motivation by the mouse and a complete halt in licking. To prevent that, we introduced a bias in the sound presentation by the target loudspeaker, increasing the likelihood of its activation to 33% at the beginning of the training and to 25% or even 17% towards the end of training, depending on the animal and its intrinsic motivation.

The scripts to control the behavioral task were custom-written in MATLAB (Mathworks).

### Data analysis and visualization

All subsequent data processing steps, calculations and statistical analyses were performed with built-in and custom-written functions in MATLAB (version R2019b).

### Data pre-processing and regions of interest (ROIs)

Raw image sequences were first corrected for phase mismatches that occur with bidirectional resonant scanning. Phase-corrected sequences were then motion-corrected using a discrete Fourier transform algorithm (Guizar-Sicairos et al., 2008). Due to the slow kinetics of GCamp6s, image sequences were then averaged by a factor of three or five to increase the signal-to-noise ratio.

ROIs were manually drawn to cover the extent of the cell body with the help of custom-written applications. This included a mask of the surrounding neuropil that extended from 3–72 pixels away from the drawn ROI (but excluded pixels pertaining to any other ROI). ROIs were excluded from analysis for sessions in which it could not be visually identified (re-found). Because of the synapsin promotor incorporated into the genomic viral expression sequence, only neurons should express the construct. However, it cannot distinguish excitatory from inhibitory neurons. Although interneurons could to some extent be manually identified by cell body size and shape, this is by no means reliable, and ROIs were thus agnostically drawn for all cell bodies that could be visually identified. Time-series from each ROI and neuropil mask were then extracted, and a factor of 0.1× the green (GCamp6s) neuropil signal was subtracted from the corresponding green ROI signal. Finally, a ratio of (neuropil-corrected) green to red (mRuby2) ROI time-series was performed in order to achieve pseudo ratiometric imaging (G/R). In total, 1228 ROIs across the twelve regions and eight were identified (though not all were re-found on each session).

For imaging experiments during the sound localization task, the time series of the fluorescence signal in the 20 µm around the ROI (excluding pixels pertaining to other ROIs) multiplied by an empirically determined factor (Kerlin et al., 2010) was subtracted from the time series of the GCamp6s fluorescence of the ROI. Large fluctuations in the fluorescence time series were removed by subtracting the 8^th^ percentile of the fluorescence measured in the time windows of 15s around each timepoint.

### Trial and ROI exclusion criteria

Strict trial exclusion criteria were applied to individual trials in order to ensure the interpretability of calcium responses. First, trials were excluded if the motion correction algorithm detected a shift between consecutive frames of more than 10 pixels (∼5 µm), which from our experience indicates a movement in *z*, for which the algorithm cannot correct. Second, for the passive experiments trials were excluded when the baseline (R_0_, 1 s before stimulus onset) exceeded 150% of the average of all trial-wise R_0_ values of that ROI in the session (Fig. 1d, top). This is because higher baseline values indicate preceding activation of GCamp6s, and due to the extreme non-linearity of calcium responses, the calcium signal after the stimulus onset in these cases could not be interpreted confidently. Third, for the passive experiments, trials were excluded if the baseline SD exceeded 0.1 times the R_0_, which indicated that the baseline was unstable or particularly noisy (Fig. 1d, top). All remaining accepted trials were then averaged and baseline-subtracted.

For the passive experiments, calcium responses (ΔR/R_0_) were calculated by taking the peak response (positive or negative) in a time window of 0.5–2 s after stimulus onset. In order to streamline subsequent statistical analyses, the values of the individual raw datapoints at the time of the peak of the averaged response were run through Kolmogorov–Smirnov (KS) tests of normality, and only those ROIs that passed the test (*P* < 0.05) were included. ΔR/R_0_ data (traces and spatial tuning functions) are thus always presented as mean ± s.e.m of the values used for the KS tests, and unless otherwise noted, only parametric statistical tests were performed on the data. Only ROIs that passed all of these exclusion criteria were deemed as valid for further analysis (valid, Extended Data Fig. 2b).

### Criteria for responsiveness and spatial tuning

Because the AC is notorious for extremely sparse responsiveness, a responsiveness threshold of 7× the s.d. (*P* ⟶ 0) of R_0_ was applied, for the passive experiments. A cell’s response for at least one loudspeaker had to equal or exceed this threshold in order to be deemed responsive. Otherwise, the cell was classified as non-responsive (NR).

In order to determine how specifically a responsive ROI was tuned to space on any given session, one-way analysis of variance (ANOVA, MATLAB function *anova1*) tests were run on all accepted trials, grouped by loudspeaker, where the best loudspeaker was defined as that which yielded the highest response. If a tuning curve passed the ANOVA (*P* < 0.05), and the best loudspeaker won comparisons (MATLAB function *multcompare*) with all other loudspeakers (*P* < 0.05), it was classified as spatial tuning specificity category (spec cat) three (Supplementary Fig. 1, right). A tuning curve that passed the ANOVA test, and whose best loudspeaker won comparisons with all loudspeakers apart from its neighbor(s), fell into spec cat two (Supplementary Fig. 1, second from right). A tuning curve that passed the ANOVA test, but did not win comparisons with all the non-neighboring loudspeakers, fell into spec cat one. (Supplementary Fig. 1, second from left; this example tuning curve would have been designated spec cat three if it had won the comparison between the zero and –60 ° loudspeakers.) Finally, any tuning curve that failed the ANOVA test was deemed non-tuned (NT, Supplementary Fig. 1, left). By definition all non-tuned neurons were responsive.

For the sound localization experiments, ΔF/F_0_ traces were determined for each neuron and loudspeaker as the mean of the green fluorescence timeseries for each timepoint across of all the trials that corresponded to that loudspeaker. The corresponding 95% confidence intervals were calculated based on bootstrapping. F_0_ was calculated per neuron as the mean of the baseline (1s before sound onset) across all trials from all loudspeakers.

For the sound localization experiments, a neuron was considered responsive if two conditions were met: (1) the mean response of the baseline (1s before sound onset) and the mean response to the sound presentation (0.1 to 1.5 s after sound onset) were significantly different (p<0.05, paired t-test) for at least one loudspeaker; (2) the maximum of ΔF/F_0_ for the significant loudspeaker was larger than the maximum of the 97.5 percentile during baseline or the minimum of ΔF/F_0_ was smaller than the minimum of the 2.5 percentile during the baseline.

For the sound localization experiments, a neuron was considered spatially tuned if: (1) it was responsive, (2) the ANOVA test was significant (p<0.05), and (3) any pairwise comparison (post-hoc Tukey-HSD test) showed significance (p<0.05).The best loudspeaker was assigned to each neuron as the loudspeaker that elicited the strongest mean response (either increase or decrease) during the sound presentation analysis period (0.1 to 1.5 s after sound onset). These best loudspeakers are represented in the flow diagram (Fig. 6e).

### Determination of data set size

For analyses of the first anesthetized and awake sessions (Figs. 1–2 and Extended Data Fig. 1b,c), all valid ROIs (in both wakefulness states) across all regions were pooled (n = 1030 of 1228 total). When only responsive ROIs between states were compared (Extended Data Fig. 1a) the subset of ROIs that were deemed responsive in both states were used (*n* = 794). Analyses of spatial tuning subdivided the respective responsive ROIs (anesthetized: *n* = 820, awake: *n* = 954) into non-tuned (NT, anesthetized: *n* = 542, awake: *n* = 612) and tuned (spec cat ≥ 1; anesthetized: *n* = 279, awake: *n* = 342).

For longitudinal analyses of passive listening experiments (Figs. 3–4), the base data set included only ROIs that were valid on all eight sessions for white noise (*n* = 698). For analyses requiring spatially tuned (Fig. 3a–d) cells, session-wise cell counts varied naturally between sessions. Visualizations of tuning switching behavior (Fig. 4b,d,e) include only cells that were spatially tuned on at least one session (awake or anesthetized, *n* = 518).

### Amplitude tuning metrics and statistics

For amplitude analyses (Extended Data Fig. 1 and Extended Data Fig. 2c) and reported amplitudes of all non-tuned cells, the maximum amplitude of all loudspeakers for each ROI was taken, regardless of the best loudspeaker on that session. The state preference indices (*I_pref_*, Extended Data Fig. 1c) were calculated with the equation:

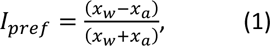

where *x_w_* and *x_a_* are corresponding values in the awake and anesthetized states, respectively. Values greater than and less than zero thus represent relatively larger values in the awake and anesthetized states, respectively. Negative values (shaded areas in Extended Data Fig. 1a), indicate a response sign inversion (values > 1 and < –1). However, because this equation has an additional inversion point at the orthogonal to the unity line (brown shaded area in Extended Data Fig. 1a), data points (*n* = 7 ROIs) in this area were excluded from the index calculations.

Although all state-wise distributions (Extended Data Fig. 1b) were deemed normal, they at times also exhibited substantial skewedness. We therefore applied the non-parametric two-sided Wilcoxon rank-sum tests of differences of medians (MATLAB function *ranksum*, Extended Data Fig. 1b) on state-wise distributions and two-sided Wilcoxon signed-rank tests (MATLAB function *signrank*, Extended Data Fig. 1c) on state preference indices.

### Pair-wise and population statistics (awake vs anesthetized)

For all pair-wise comparisons of individual ROI response amplitudes between wakefulness states (Extended Data Fig. 1a), one-way ANOVAs were performed on the valid trials in each state, and a threshold was set at *P* < 0.05. To evaluate statistically significant differences in spatial tuning functions between states (Fig. 3b), a two-way ANOVA (MATLAB function *anovan*) was performed on the response amplitudes to each loudspeaker and state and evaluated for statistical significance between speakers and states.

### Longitudinal analyses (awake vs anesthetized)

Because the amplitude responses in the longitudinal datasets were not normally distributed, quantile normalization was first performed (MATLAB function *quantilenorm*) to normalize the statistics of each statistical group (i.e. session 1, awake). To quantify factorized changes in amplitude through each state transition (7 in total, Fig. 4b), the amplitudes after the transition were divided by the amplitudes observed before. To evaluate differences in spatial tuning between states and over sessions, these data were run through a three-way ANOVA (Fig. 4b). To evaluate the statistical significance of tuning changes over sessions for each ROI independently (Fig. 4f), two way-ANOVAs were performed (on all or just awake and anesthetized sessions) and neurons were subdivided by their corresponding *P* values (threshold: 0.05).

### Behavioral data analysis

To analyze both the simultaneous behavior and imaging data, we developed custom written MATLAB (Mathworks) and Python scripts. The performance of the mice was represented as the lick rate in response to the presentation of sounds by each loudspeaker. The associated 95% confidence interval was calculated with bootstrapping. To determine that the mouse licked significantly more to the rewarded loudspeaker than to another, the Bonferroni corrected confidence interval (divided by the six loudspeakers – corresponding to the 99.17% confidence interval) to the rewarded loudspeaker could not intercept the real lick rate to the non-rewarded loudspeaker and vice-versa, the real lick rate to the rewarded loudspeaker could not intercept the Bonferroni corrected confidence interval to the non-rewarded loudspeaker. To choose the sessions used for the analysis, we compared the number of neurons found in all sessions before and after learning. The sessions with the highest number of common neurons were then selected.

## Acknowledgments

This work was supported by the Max Planck Society (MPI Fellow Research Group to B.G.) and the German Research Foundation (DFG) via the Munich Cluster for Systems Neurology (SyNergy; B.G.) and the DFG Research Training Group (RTG) 2175 Perception in Context and its Neural Basis (C.L. and B.G.; Stipend to M.G.). We thank Julia Kuhl (Max Planck Institute for Biological Intelligence) for help with a schematic illustration.

## Figure captions

**Extended Data Fig. 1: Neuronal responses are larger in the anesthetized than in the awake state. a**, Scatter plot of response amplitudes in the first awake versus anesthetized states (for cells deemed responsive in both states). Filled markers indicate ROIs that had statistically significant response amplitude differences between states (*P* < 0.05, one-way ANOVA). **b**, Histograms of calcium response amplitudes of all (analysis-valid) cells in the first anesthetized (blue) and awake (red) sessions, with the number of non-responsive cells in each state shown on the left. Solid lines indicate the corresponding median values, and the *P* value is the result of a Wilcoxon rank-sum test. **c**, Histogram of a calculated state preference index, where cells with values < –1 and > 1 represent those in the corresponding shaded areas in **a** (sign inversion). Cells in the brown highlighted area in **a** were excluded from the index, the solid black line indicates the median of the population, and the *P* value is the result of a Wilcoxon signed-rank test.

**Extended Data Fig. 2: General response and tuning metrics of the longitudinal dataset. a**, Cut-outs of individual example ROIs across all eight sessions showing the general quality of re-finding cells from session to session. Color channels are the same as in Figure 1c,d, but are normalized to the maximum fluorescence of the structural marker (mRuby2) for each ROI and session. **b**, General overview of the number of cells valid for analysis (valid, circles), responsive cells (resp., squares) and tuned cells (tune., triangles) across sessions. Data are separated by evaluating each session individually (any, filled markers) and taking the cumulative number across sessions (cum., empty markers). Evaluation over all sessions regardless of state are shown on the left, with that of only anesthetized and awake sessions shown in the middle and on the right, respectively. The upward pointing arrow shows which data point corresponds to the number of cells that were valid (*n* = 698) on all sessions. **c** Cumulative density functions (top) and box plots (bottom) of response amplitudes of all sessions. *P* values are the results of a two-way ANOVA after quantile normalization.

**Extended Data Fig. 3: Neuron counts of tuning categorization for both FOVs in all mice over all sessions.** FOVs, and mice are organized into rows and columns, respectively. Anesthetized and awake sessions are color-coded according to the color scale at the bottom right panel, and *n* values correspond to the number of valid ROIs (over all sessions) in each FOV.

**Supplementary Fig. 1:** Example spatial tuning functions showing the determination and categorization of spatial tuning specificity (spec cat). Arrows indicate the best loudspeaker.

