## Supplementary figures and images for "The Dynamics of Context-Dependent Space Representations in Auditory Cortex"

### Extended and supplementary data and figures

#### Extended Data Fig. 1

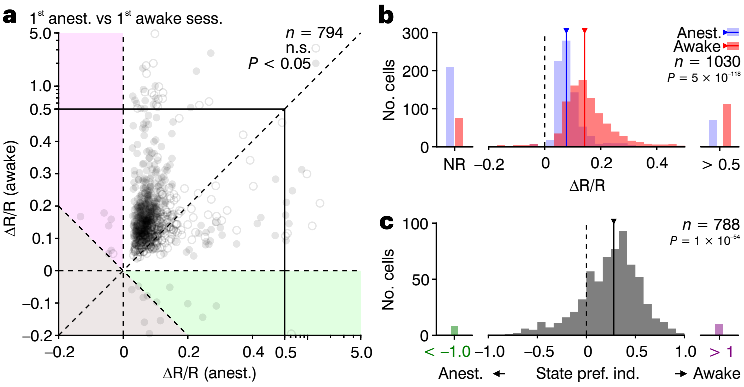

#### Extended Data Fig. 2

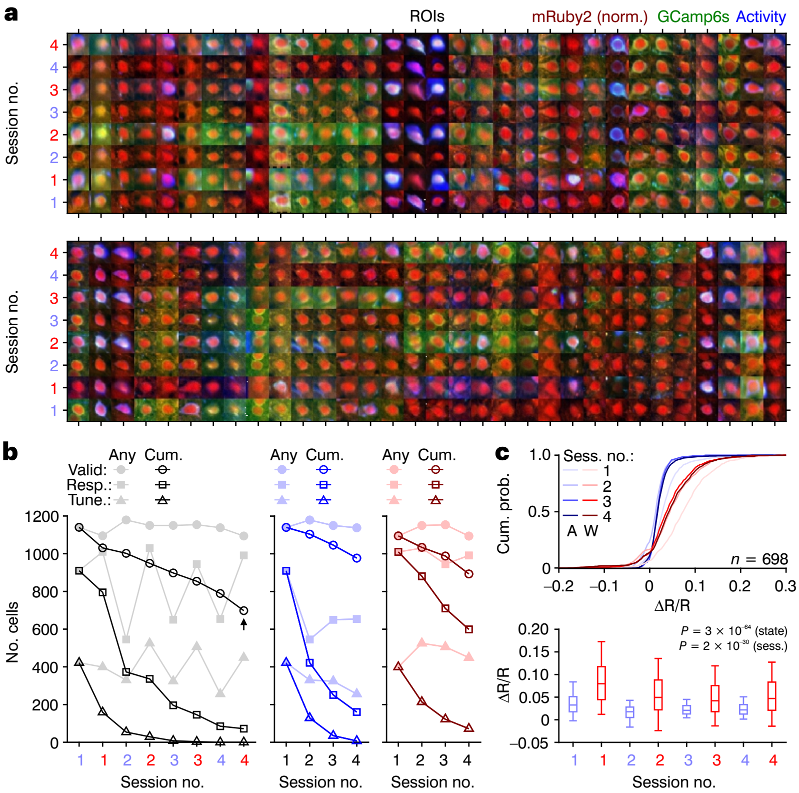

#### Extended Data Fig. 3

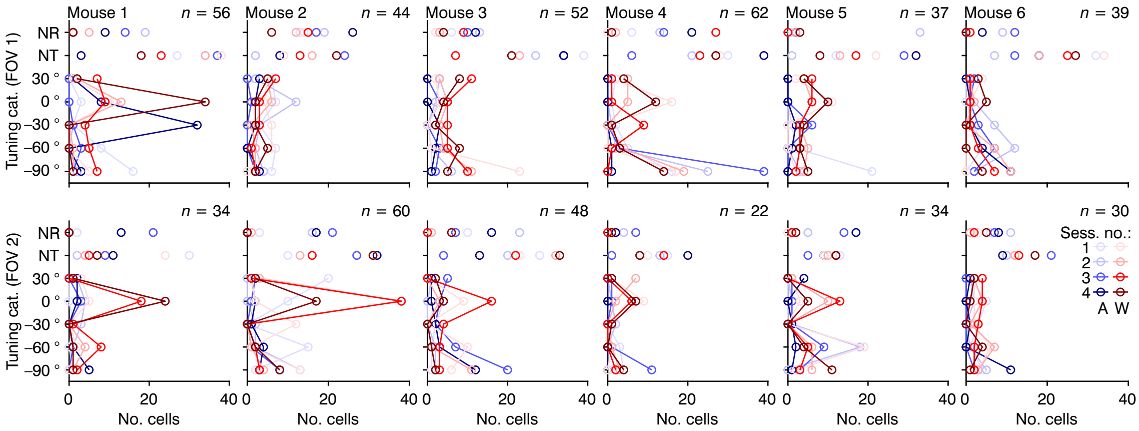

#### Supplementary Fig. 1

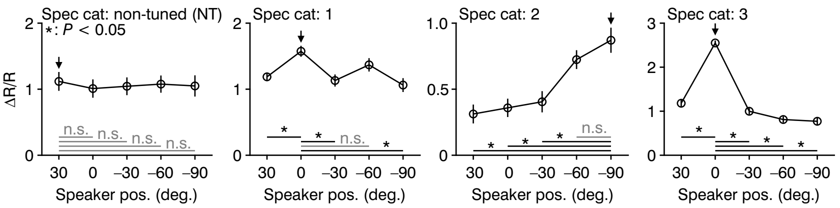
